# Identification of a role for an *E6-like 1* gene in early pollen-stigma interactions in *Arabidopsis thaliana*

**DOI:** 10.1101/623686

**Authors:** Jennifer Doucet, Christina Truong, Elizabeth Frank-Webb, Hyun Kyung Lee, Anna Daneva, Zhen Gao, Moritz K. Nowack, Daphne R. Goring

**Author notes:** Corresponding author: Daphne Goring, E-mail address.

## Abstract

In *Arabidopsis*, successful pollen-stigma interactions are dependent on rapid recognition of compatible pollen by the stigmatic papillae located on the surface of the pistil and the subsequent regulation of pollen hydration and germination, and followed by the growth of pollen tubes through the stigma surface. Here we have described the function of a novel gene, *E6-like 1* (*E6L1*), that was identified through the analysis of transcriptome datasets, as one of highest-expressed genes in the stigma, and furthermore, its expression was largely restricted to the stigma and trichomes. The first *E6* gene was initially identified as a highly-expressed gene during cotton fiber development, and related E6-like predicted proteins are found throughout the Angiosperms. To date, no orthologous genes have been assigned a biological function. Both the *Arabidopsis* E6L1 and cotton E6 proteins are predicted to be secreted, and this was confirmed using an E6L1:RFP fusion construct. To further investigate *E6L1*’s function, one T-DNA and two independent CRISPR-generated mutants were analyzed for compatible pollen-stigma interactions, and pollen hydration, pollen adhesion and seed set were mildly impaired for the *e6l1* mutants. This work identifies *E6L1* as a novel stigmatic factor that plays a role during the early post-pollination stages in *Arabidopsis*.

**Key Message:** We describe a function for a novel *Arabidopsis* gene, *E6-like 1* (*E6L1*), that was identified as a highly-expressed gene in the stigma and plays a role in early post-pollination stages.

## Introduction

The characteristic dry stigmas of the Brassicaceae lack surface secretions, that can enable spontaneous pollen hydration and germination, and this trait allows the stigmas to regulate the early post-pollination stages. Following a compatible pollination, several physiological steps follow, starting from pollen adhesion, pollen hydration, pollen germination, and resulting in pollen tube entry into the stigma. The contact zone, between a pollen grain and a stigmatic papilla, is where the pollen recognition is initiated and is composed of proteins and lipids from the pollen coat and papillar surface (reviewed in Doucet et al. 2016; Goring 2017; Zheng et al. 2018). It is also at this stage that self-incompatible pollen is recognized and rapidly rejected by blocking the physiological steps which begin at pollen hydration (reviewed in Doucet et al. 2016; Goring 2017; Jany et al. 2019). A number of players have been identified to be required for compatible pollen-stigma interactions, and they broadly fall into three categories.

The first group includes mutants with altered structural components that will in turn alter post-pollination processes (reviewed in Zheng et al. 2018). For example, the *Arabidopsis eceriferum* (*cer*) mutants have defects in lipid biogenesis that causes a loss of the pollen coat and impaired pollen hydration and germination (Preuss et al. 1993; reviewed in Jiang et al. 2013). In contrast to this phenotype, the *Arabidopsis fiddlehead* (*fdh*) mutant has altered cuticle biogenesis which allows for the germination of wild-type pollen on non-reproductive plant tissues (Lolle and Cheung 1993; Voisin et al. 2009). Increased cell-wall permeability was also observed in the *fdh* mutant and was proposed as an explanation for this phenotype (Lolle et al. 1997). Cell wall modifications are also important on both sides since the *Arabidopsis* pollen *O-fucosyltransferase1* (OFT1) was observed to be required for pollen tube penetration through the stigma and style (Smith et al. 2018a; Smith et al. 2018b). Moreover, modifications of cell wall components in the transmitting tract are required to allow for pollen tube growth and travel (reviewed in Herrera-Ubaldo and de Folter 2018).

The second group is represented by proteins involved in the perception of compatible pollen. In early studies, two secreted *Brassica* glycoproteins, SLR1 and SLG, were implicated in the stigma for pollen adhesion (Luu et al. 1997; Luu et al. 1999). Interestingly, both of these secreted glycoproteins were also demonstrated to bind to small *Brassica* class-A pollen coat proteins (PCPs; Doughty et al. 1993; Hiscock et al. 1995; Takayama et al. 2000). More recently, class-B PCPs were demonstrated to be required for pollen adhesion and hydration on stigmas as pollen grains from a triple *pcp-b* mutant were impaired in these processes (Wang et al. 2017). The final group is comprised of players connected to cellular responses required to promote the physiological steps following compatible pollen recognition. On the pollen side, a role for ROS production has been implicated for pollen hydration and germinating pollen tubes in studies using mutant pollen grains from *Arabidopsis rboh* (NADPH oxidase; Kaya et al. 2015) and *Arabidopsis kinβγ* (Gao et al. 2016; Zheng et al. 2018) mutants. In addition, the function of *Arabidopsis* KINβγ was connected to the expression of an inward shaker K^+^ channel gene, *SPIK*, which again impaired pollen hydration when mutated (Li et al. 2017). Further down, the endomembrane trafficking protein, VPS41, was shown to be required in *Arabidopsis* pollen for pollen tube penetration through the stigma into the transmitting tract (Hao et al. 2016). On the stigma side, components associated with vesicle trafficking, such as the exocyst complex and phospholipase Dα1, are required in *Arabidopsis* and *Brassica* for pollen-stigma interactions (Safavian and Goring 2013; Safavian et al. 2014; Safavian et al. 2015; Samuel et al. 2009; Scandola and Samuel 2019). The ACA13 Ca^2+^ ATPase is also required in the stigma for compatible pollen acceptance (Iwano et al. 2014).

In this study, we investigate the function of a potential new player in pollen-stigma interactions: a novel gene that is highly-expressed in the stigma and annotated as an *E6-like protein* (Araport11). This annotation is based on its predicted sequence similarity to the cotton *E6* gene which was first identified as a highly-expressed gene in cotton fibers (Ji et al. 2003; John 1996; John and Crow 1992; Li et al. 2002). Cotton fibers are single cell trichomes found on the cotton ovule epidermal surface and their development is characterized by several distinct phases including fiber initiation, elongation, secondary cell wall synthesis and maturation. The elongation phase is carried out by rapid growth without cell division, and is considered an example of polarized cell growth analogous to leaf trichome development (reviewed in Lee et al. 2007). Cotton *E6* gene expression was found to be upregulated in mature ovules, with levels highest at the elongation phase of the cotton fibers, and absent from other tissues such as stems and leaves (Indrais et al. 2011; Lee et al. 2006; Lee et al. 2007). Despite the high expression associated with cotton fibers, there are no known functions for cotton *E6* (or other plant *E6-like* genes). Antisense *E6* knockdown lines in *Gossypium hirsutum* did not display any phenotypic changes for cotton fiber development, and the cotton fiber properties remained unaffected (John 1996). Here, we explore the function of the *Arabidopsis E6-like protein* gene, now referred to as *E6-like-1* (*E6L1*), in the stigma following compatible pollinations through the analysis of T-DNA and CRISPR-knockout *e6l1* mutants.

## Materials and Methods

### Plant Materials and Growth Conditions

*Arabidopsis* seeds were sterilized, stratified at 4°C, and then transferred directly to soil or first plated on ½ Murashige and Skoog (MS) medium plates for 7-10 days and then transferred to soil. To view trichomes, seedlings were germinated and grown on ½ MS plates with 1% sucrose for two weeks. *Nicotiana benthamiana* seeds were cold stratified for several days and planted directly on soil. All soil was supplemented with 1g/L 20-20-20 fertilizer, and plants were grown at 22°C on a 16h light/8 h dark cycle. Humidity was monitored in the growth chambers and maintained at between 20-60% relative humidity.

### Expression Profiling and Phylogenetic Analysis

To identify novel genes showing stigma enriched expression, the BAR Expression Angler tool (http://bar.utoronto.ca/; Toufighi et al. 2005) was used with the stigma-specific expressed SLR1 gene as the bait (At3g12000; Dwyer et al. 1992) and the *AtGenExpress Plus-Extended Tissue Compendium* (Schmid et al. 2005) as the dataset being searched. Stigma/ovule (Swanson et al. 2005) and trichome (Gilding and Marks 2010; Marks et al. 2009) microarray datasets were accessed through the BAR. Additional datasets used were the stigmatic papillae RNA-Seq dataset (Gao et al. 2018), stigma microarray datasets (Iwano et al. 2014), and the TRAVA RNA-Seq dataset (http://travadb.org/; Klepikova et al. 2016). Expression profiling data was displayed using the HeatMapper Plus tool or Arabidopsis eFP browser tool at the BAR (Toufighi et al. 2005; Winter et al. 2007). For the *Arabidopsis* E6-like genes, gene annotations and amino acid sequences were retrieved from TAIR and Araport11 (Berardini et al. 2015; Krishnakumar et al. 2015).

For the phylogenetic analysis of angiosperm E6-like proteins, protein BLAST searches (Altschul et al. 1997) were performed in *EnsemblPlants* (Kersey et al. 2018), *Phytozome* (Goodstein et al. 2012), *GymnoPLAZA* (Proost et al. 2015), NCBI (Coordinators 2017) and the *Brassica* Database (Wang et al. 2015). Amino acid sequences were retrieved from *EnsemblPlants* (*Actinidia chinensis, Beta vulgaris, Arabidopsis lyrata, Brassica oleracea, Corchorus capsularis, Cucumis sativus, Daucus carota, Medicago truncatula, Phaseolus vulgaris, Gossypium raimondii, Prunus persica, Nicotiana attenuata, Solanum lycopersicum, Theobroma cacao, Vitis vinifera, Brachypodium distachyon, Leersia perrieri, Oryza sativa Japonica, Hordeum vulgare, Setaria italica, Sorghum bicolor, Amborella trichopoda*); *Phytozome* (*Aquilegia coerulea, Gossypium raimondii, Citrus clementina*); NCBI (*Tarenaya hassleriana*) and the *Brassica* Database (*Aethionema arabicum*) (File S1). The MEGA7 software (Kumar et al. 2016) was used to produce multiple protein sequence alignments using Muscle (Edgar 2004). The predicted N-terminal signal peptides were trimmed in the Muscle alignment (File S2) and then used to generate a consensus tree by the Maximum Likelihood method (Jones et al. 1992) with 1000 bootstrap replicates (Felsenstein 1985) in MEGA7. Additional alignments were created in MEGA7 (Files S3-S7) to use in the WEBLOGO tool for generating consensus sequence logos (https://weblogo.berkeley.edu/logo.cgi; Crooks et al. 2004).

### Vector construction and plant transformation

The *E6L1* cDNA was amplified by RT-PCR from Col-0 stage 12 whole bud RNA using Taq polymerase (see Table S6 for primers) and cloned into the TOPO entry clone using the PCR8/GW TOPO cloning kit (ThermoFisher Scientific). Gateway reactions were carried out using LR clonase II enzyme (ThermoFisher) to transfer the *E6L1* cDNA (minus stop version) into the pUBC-RFP-Dest vector and generate a C-terminal *E6L1:RFP* fusion under the control of the *UBQ10* promoter (Grefen et al. 2010). The *E6L1:RFP* vector was then electroporated into *Agrobacterium* GV2260 for the agroinfiltration experiments. Leaves 3 or 4 from 5-week-old *N. benthamiana* leaves were then transformed with *E6L1:RFP* by agroinfiltration as described by (Sparkes et al. 2006).

Three vectors were obtained for the generation of CRISPR/Cas9 vectors: pHEE401E and pBUE411 from Addgene, and pCBC-DT1T2 from ABRC (Wang et al. 2015; Xing et al. 2014). pHEE401E contains a *HygR* selectable marker in the T-DNA, and a version was generated that carried the *BlpR* selectable marker instead using the pBUE411 vector as a source of the *BlpR* gene. Both plasmids were digested with EcoRI/MreI which dropped out a ~3.9 kb fragment from pBUE411 containing *35S:BlpR* and the corresponding fragment from pHEE401E containing *35S:HygR*. The ~3.9 kb EcoRI/MreI *35S:BlpR* containing fragment was then ligated to the ~12 kb EcoRI/MreI pHEE401E vector backbone fragment to generate pBEE401E. All CRISPR/Cas9 vectors targeting the *E6L1* gene were then generated using pBEE401E. The CRISPR sgRNA (single guide RNA) sequences targeting *E6L1* were selected using the CHOPCHOP software to search for sequences adjacent to PAM sites and avoid potential off-targets in the *A. thaliana* genome (Labun et al. 2016). A PCR fragment containing two sgRNA target sequences was generated using the pCBC-DT1T2 vector template with the Phusion polymerase (NEB; See Table S6 for primers), and the purified PCR fragment was then cloned into pBEE401E using a golden gate reaction with BsaI (Wang et al. 2015; Xing et al. 2014). Two pBEE401E-based vectors carrying either the CR03-CR01 or CR04-CR01 sgRNA combinations were transformed into *Agrobacterium* by electroporation, and *Arabidopsis* plants were then transformed using the floral dip method (Clough and Bent 1998). Seeds collected from the dipped plants were stratified and planted directly on soil, and flats of seedlings were screened for transgenic plants by spraying with the Basta herbicide.

### Generation of *e6l1* mutants

The *Arabidopsis e6l1-1* T-DNA line (GK-763B03) was obtained from ABRC, and homozygous mutations were confirmed by PCR. The location of the T-DNA was also verified by sequencing the genotyping PCR products. The *e6l1-2* and *e6l1-3* mutant alleles were generated with the CRISPR constructs described above. Deletions generated in the *E6L1* gene were verified using primers flanking the sgRNA target regions. Two independent T1 lines were isolated from the two constructs (CR03-CR01 or CR04-CR01) and brought to the T2 and T3 generations to isolate homozygous mutants (*e6l1-2* and *e6l1-3*). Additionally, a set of primers within the deleted region were used to verify that the *e6l1* mutants were indeed homozygotes (where a PCR product could be obtained for those plants carrying the wild-type *E6L1* allele while no PCR product was observed for the homozygous deletion plants). See Table S6 for primers.

### Assays for pollen hydration, pollen adhesion and pollen tube growth, and seed set

All pollination assays were conducted as previously described and performed under an ambient humidity of less than 60% (Safavian et al. 2014; Safavian et al. 2015). Briefly, stage 12 flower buds were emasculated and carefully wrapped with plastic wrap and allowed to mature overnight. For pollen hydration assays, pistils were mounted upright in ½ MS medium and hand-pollinated with a small amount of Col-0 pollen. Pictures were taken immediately at 0 min and again at 10 min post-pollination using a Nikon sMz800 stereo zoom microscope at 6.3× magnification with a 1.5× objective. Pollen grain diameter was measured laterally using the Nikon digital imaging software for 10 random pollen grains per pistil, 3 pistils per genotype. For pollen adhesion and pollen tube growth, pistils were lightly pollinated with pollen and then collected at 2h post-pollination, fixed and stained with aniline blue. Pollinated pistils were imaged using a Zeiss Axioskop2Plus microscope under bright-field to count the number of pollen grains adhered, and under UV fluorescence to image pollen tube growth. Pollen adhesion was quantified 10 pistils for each cross. For counting seed set, green siliques were collected from naturally self-pollinated flowers and sliced longitudinally to count the number of developing seeds. 10 siliques were counted for each pollination.

### Microscopy

For confocal microscopy, leaf disks were cut from *N. benthamiana* at 48 hours post-infiltration and visualized using a Leica TCS SP5 confocal microscope. Image processing was done using the Leica LAS AF lite software. Plasmolysis was achieved by treatment with 0.8M mannitol as described by Lang *et al.* (2014).

For scanning electron microscopy, each individual pistil sample was collected from a freshly-opened stage 13 flower (from either Col-0, *e6l1-1*, *e6l1-2* or *e6l1-3* plants) and immediately mounted on top of conductive tape for imaging. The top views of the stigmas were imaged at an accelerating voltage of 3kV and a magnification of ×200. All images were taken by using the Hitachi SU3500 scanning electron microscope. Image J (Schneider et al. 2012) was used on the SEM images to measure angles of deviation for stigmatic papillae.

The Nikon sMz800 stereo zoom microscope was used to photograph trichomes, stigmas and siliques.

## Results

### The *Arabidopsis E6L1* gene is highly expressed in the stigma

To identify potential new candidate genes involved in pollen-stigma interactions, we performed a search for genes displaying enriched expression in the stigma. This was carried out by using the BAR *Expression Angler* tool (Toufighi et al. 2005) with the stigma-specific expressed SLR1 gene as the bait (At3g12000; Dwyer et al. 1992; Franklin et al. 1996). The *Arabidopsis E6L1* gene (At2g33850) was ranked 7^th^ on this list (Fig. 1A, Table S1-S2) and became of particular interest as it was one of the top ranked genes based on expression levels in a stigmatic papillae RNA-Seq dataset (Fig. 1B, Table S3; Gao et al. 2018) and stigma microarray datasets (Table S4; Iwano et al. 2014). *E6L1* is annotated as a ‘E6-like protein’ due to its amino acid similarity to the cotton E6 protein (AAB03079.1; alignment shown in Fig. S1). There were also two other top ranked genes predicted to encode secreted proteins in the stigmatic papillae RNA-Seq dataset (*LTP1* and *PRP4*, Fig. 1B). However, *E6L1* was unique in having a largely stigma-specific expression pattern when surveyed across all the different tissues in the TRAVA RNA-Seq dataset (Fig. 1C, Table S5; Klepikova et al. 2016). In addition, there are two other *Arabidopsis E6-like* genes: At1g03820 (annotated as a ‘E6-like protein’) and At1g28400 (mis-annotated as a ‘GATA zinc finger protein’). However, these genes displayed little or no expression in the stigmatic papillae RNA-Seq dataset (Table S3, see also Table S5 for TRAVA RNA-Seq expression patterns). Thus, the high-level stigma-enriched expression pattern for *E6L1* made it an interesting candidate gene to study in the context of pollen acceptance. Furthermore, a survey of the microarray expression datasets for different tissues in the BAR eFP browser uncovered *E6L1* expression in trichomes in addition to the stigma (Fig. 1A, Table S2). This is also an interesting feature given that the cotton *E6* gene was originally identified for its high expression in cotton fibers which represent seed trichomes (John 1996; John and Crow 1992; Lee et al. 2006; Lee et al. 2007).

**Fig. 1.**
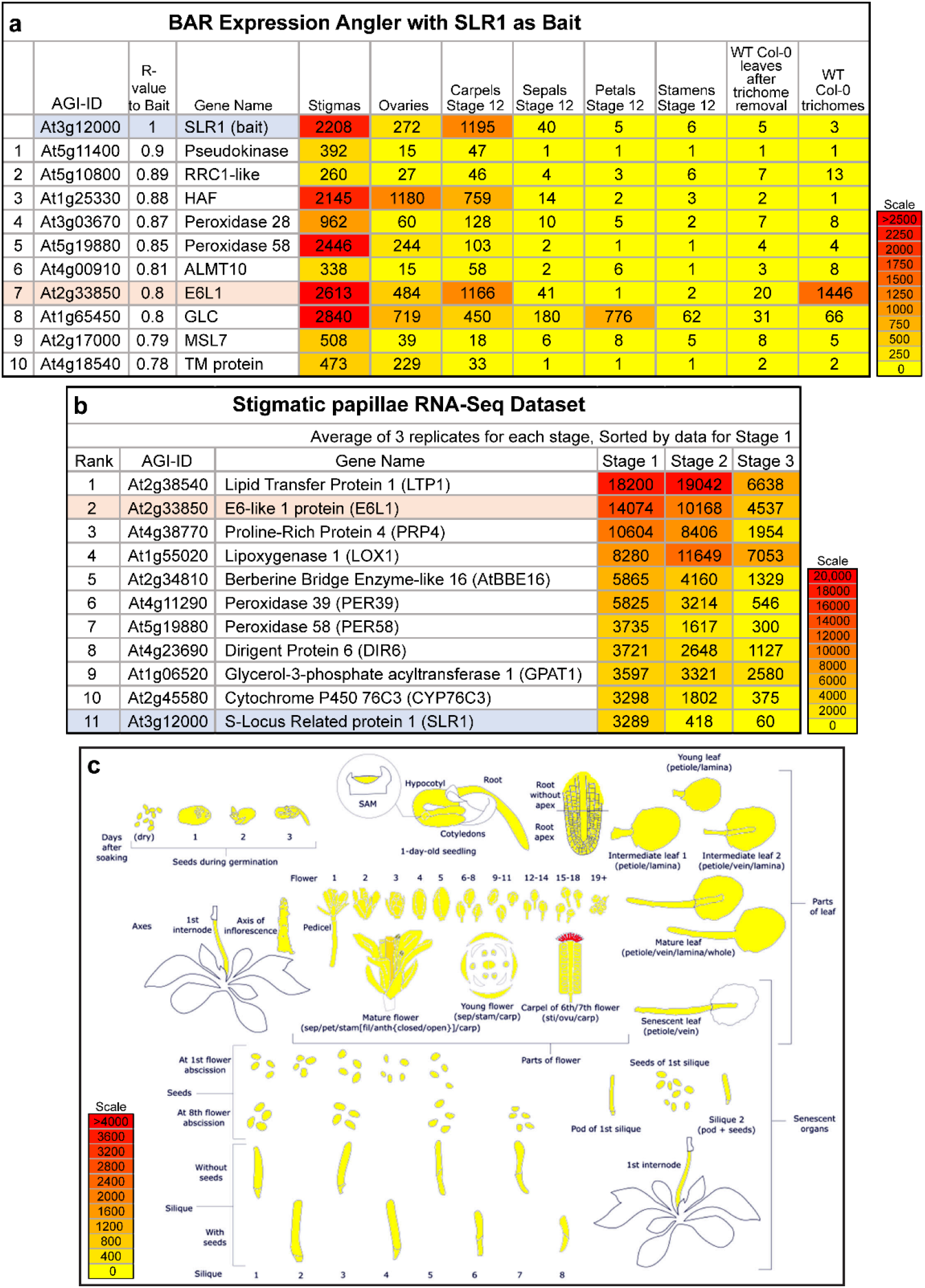
*E6L1* expression profiles in microarray and RNA-Seq datasets. **a.** BAR *Expression Angler* search for genes that display enriched expression in the stigma. The genes were identified by using the BAR *Expression Angler* tool with the stigma-specific *SLR1* gene (At3g12000) as the bait (Dwyer et al. 1992; Franklin et al. 1996; Toufighi et al. 2005). The top 10 ranked genes are shown (*SLR1* highlighted in blue, *E6L1* highlighted in orange). See Table S1-S2 for the full dataset with the top 25 ranked genes. **b.** Stigmatic papillae RNA-Seq Dataset sorted by expression levels. The top 11 ranked genes are shown, based on Stage 1 (Stage 13 flowers) expression levels (*E6L1* highlighted in orange*, SLR1* highlighted in blue). See Table S3 for the top 25 ranked genes and staging information. **c.** *E6L1* expression profile of across different tissues. High expression levels are only shown in the stigma. This data is derived from the TRAVA RNA-Seq dataset as displayed in the *Klepikova Arabidopsis Atlas eFP browser* (http://bar.utoronto.ca/; (Klepikova et al. 2016; Winter et al. 2007). See Table S5 for the relative read counts.

### The *Arabidopsis E6L1* gene encodes a secreted protein and belongs to conserved family of predicted genes encoding E6-like proteins in Angiosperms

The *E6L1* gene is predicted to encode a protein of 267 amino acids in length. Similar to the predicted cotton E6 protein, *Arabidopsis* E6L1 has an unknown function, and by using the SUBA4 and SignalP prediction tools, is predicted to be secreted to the cell wall via a signal peptide (Fig. S2; Hooper et al. 2017; Petersen et al. 2011). The other general feature of the predicted cotton E6 and *Arabidopsis* E6L1 proteins is that ~15% of the amino acids are asparagine residues and represent potential (N)-linked glycosylation sites (Fig. S1; Strasser 2016). In order to test the prediction that E6L1 is secreted, a C-terminal RFP fusion under the control of a UBQ10 promoter, was generated and transiently expressed in *N. benthamiana* leaf epidermal cells (Grefen et al. 2010; Sparkes et al. 2006). Confocal laser scanning microscopy images for E6L1:RFP displayed an RFP signal at the cell periphery which would be consistent with localization to the plasma membrane and/or cell wall (Fig. 2A-C). In order to differentiate between these two compartments, transformed cells were plasmolysed by treatment with 0.8M mannitol. Upon plasmolysis, a diffuse RFP signal was observed at the cell periphery which would be expected for a protein being secreted to the apoplastic space (Fig. 2D-F).

**Fig. 2.**
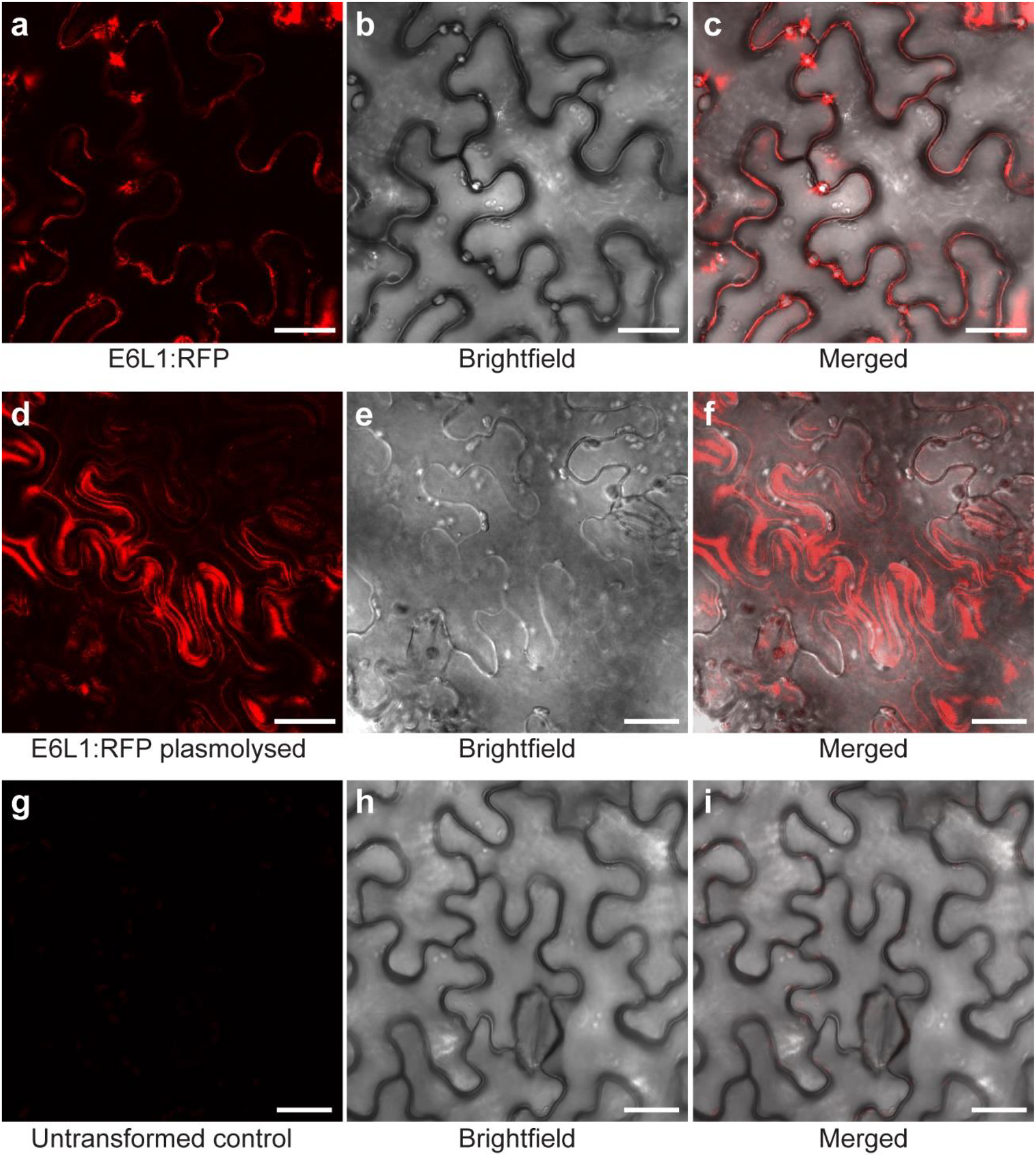
Transient expression and subcellular localization of E6L1:RFP in *N. benthamiana* leaves. **a-c.** E6L1:RFP appears to be localized around the cell periphery. **d-e.** E6L1:RFP is diffusely localized to the cell periphery and apoplastic space when cells were plasmolysed for 30 min with 0.8M mannitol. **g-i.** Untransformed control leaf displays very low levels of chloroplast autofluorescence. Agrobacteria carrying the *E6L1:RFP* construct were infiltrated into *N. benthamiana* leaves at OD_600_ = 0.5, and confocal images were taken at 48h post-inoculation. Scale bars = 25 μm.

To gain more insight into the E6-like proteins, the cotton and *Arabidopsis* amino acid sequences were used in protein BLAST searches against other plant genomes and a number of homologues were identified. In the process of aligning these amino acid sequences, one region was noticed to be highly conserved across the different predicted E6-like proteins and was used to generate a consensus sequence logo (Fig. 3a). This consensus sequence proved to be even more effective in protein BLAST searches to specifically identify homologous E6-like proteins when tested in the *EnsemblPlants*, *Phytozome* and *GymnoPLAZA* databases. From these different protein BLAST searches, one or more predicted proteins annotated as ‘Protein E6-like’ were identified in all angiosperm genomes tested, including the basal angiosperm, *Amborella trichopoda*. No homologues were identified in genomes for gymnosperms and basal land plants. This also is consistent with the InterPro “Protein E6-like’ classification that only lists homologues for angiosperm species (IPR040290, Mitchell et al. 2019). A cross-section of different angiosperm E6-like homologues were aligned, and the highly conserved region corresponding to the consensus sequence logo is shown for a subset of E6-like proteins in Fig 3b. The full alignment was also used to generate a new consensus sequence logo across the whole E6-like sequence, and a few other smaller stretches with some conservation were observed (Fig. S3). Finally, a phylogenetic tree was generated to examine relationships between the different angiosperm E6-like homologues (Fig. 4). There are four clades that are supported by bootstrap values and contain family-specific members. The first clade (at the top) contains Malvaceae E6-like homologues including the cotton E6 protein; however, there are also Malvaceae E6-like homologues found in other clades. The second clade (on the left) specifically has all the Poaceae (grasses) E6-like homologues. The final two supported clades (at the bottom) include all the Brassicaceae E6-like homologues along with the closely-related Cleomaceae species, *Tarenaya hassleriana*.

**Fig. 3.**
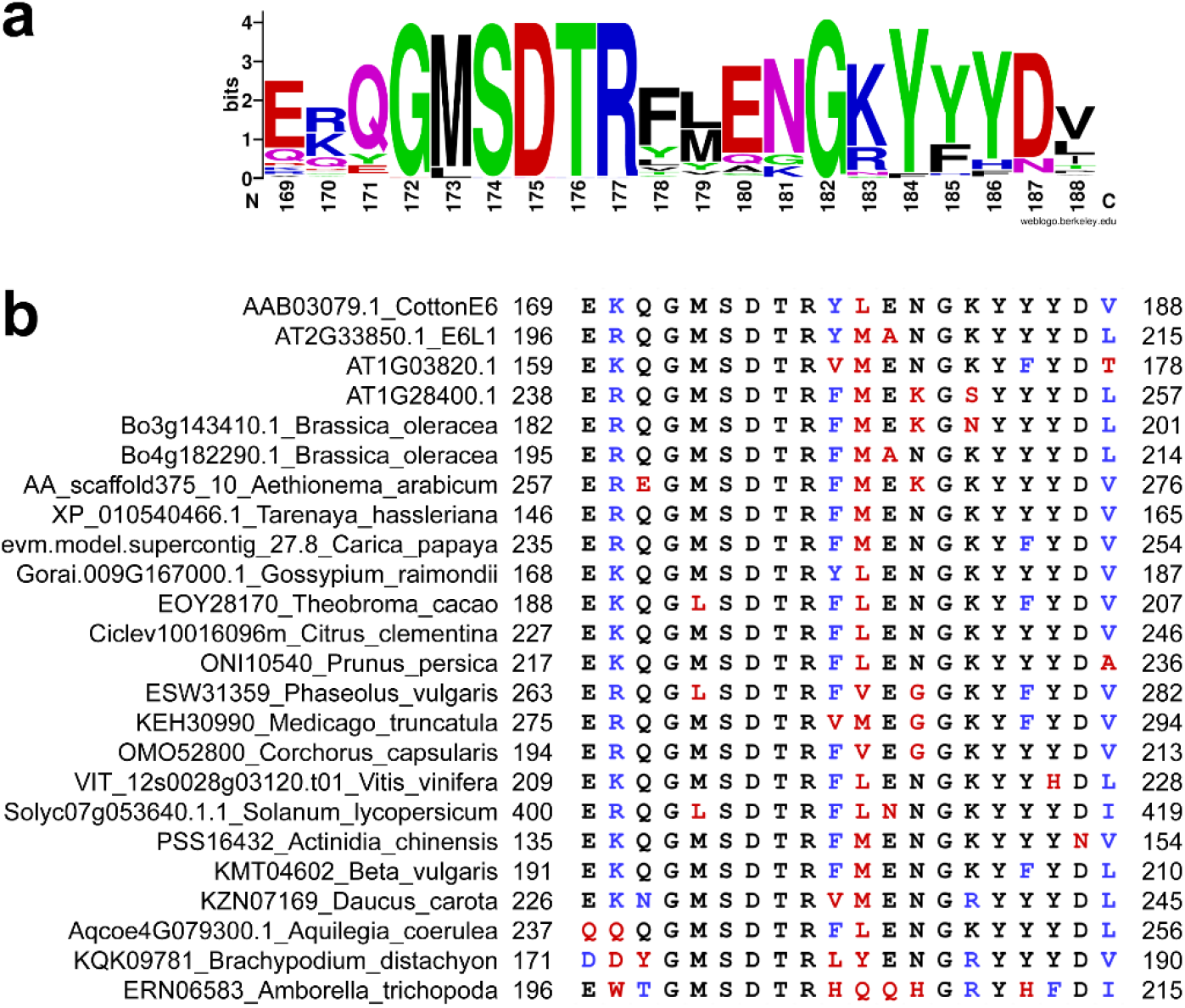
Consensus sequence for a highly conserved region in predicted angiosperm E6-like proteins. **a.** An alignment of 72 sequences for the conserved region was used to generate a Consensus Sequence Logo using the WEBLOGO tool (Crooks et al. 2004). The numbering on the X-axis represents the position of this conserved region in the cotton E6 protein sequence. **b.** Sequences for the highly conserved region are shown from 24 predicted angiosperm E6-like proteins. This represents a subset of the predicted E6-like proteins surveyed (see Files S3 and S6 for the full and conserved domain alignments, respectively).

**Fig. 4.**
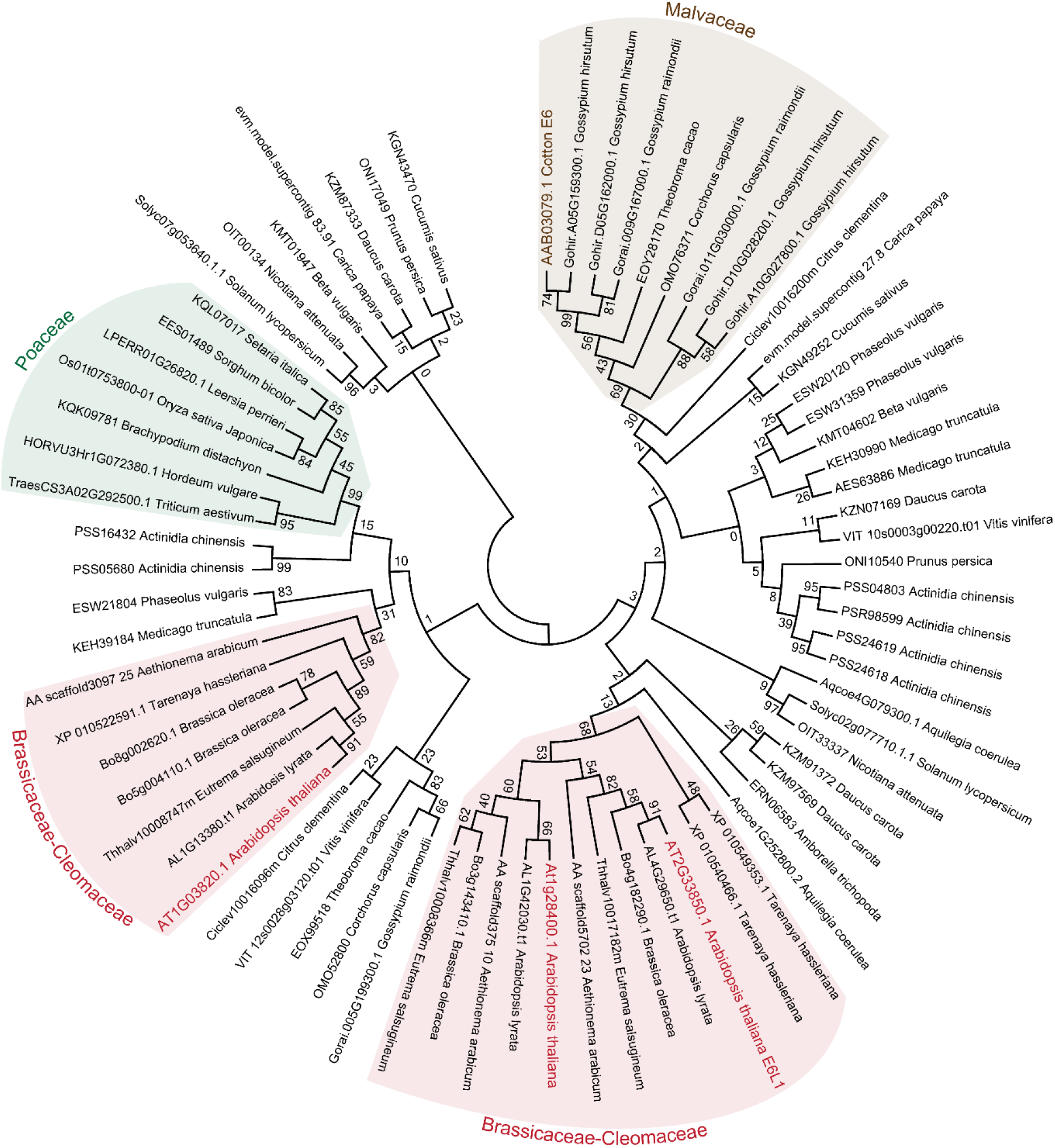
Phylogenetic analysis of predicted Angiosperm E6-like proteins. Protein BLAST searches were used to identify predicted E6-like proteins in plant genomes, and homologues were only identified in angiosperm species. From these searches, 73 amino acid sequences from a cross-section of 28 angiosperm genomes (File S1) were aligned using Muscle (Edgar 2004) in the MEGA 7 software (Kumar et al. 2016). The predicted N-terminal signal peptides were trimmed (see File S2 for alignment), and the tree was constructed using the Maximum Likelihood method in the MEGA 7 software. For the phylogenetic analysis, all positions containing gaps and missing data were eliminated, and a total of 57 positions were in the final dataset. The tree generated in MEGA 7 represents the bootstrap consensus tree inferred from 1000 replicates. See results, and materials and methods for full details. Four supported clades display family-specificity are shaded.

### *Arabidopsis E6L1* knockout mutants displayed a mild impairment in pollen-stigma interactions

In order to study the function of *E6L1* in the stigma, three independent mutants were generated and characterized for compatible pollen responses. One T-DNA mutant was obtained with a T-DNA inserted in the middle of the single exon and named *e6l1-1* (Fig. 5A). Loss of *E6L1* expression was confirmed in the homozygous *e6l1-1* mutant using RT-PCR with RNA extracted from stage 12 flower buds (Fig. 5B). Two additional *e6l1* mutant alleles were then generated with a CRISPR/Cas9 system that uses two separate sgRNAs to generate deletions (Wang et al. 2015). To produce these CRISPR *e6l1* mutants, *Arabidopsis* plants were transformed with either the CR03-CR01 or CR04-CR01 sgRNA combinations, and homozygous deletion mutants for each combination were identified in the T3 generation for further analysis (Fig 5C, *e6l1-2* and *e6l1-3*).

**Fig. 5.**
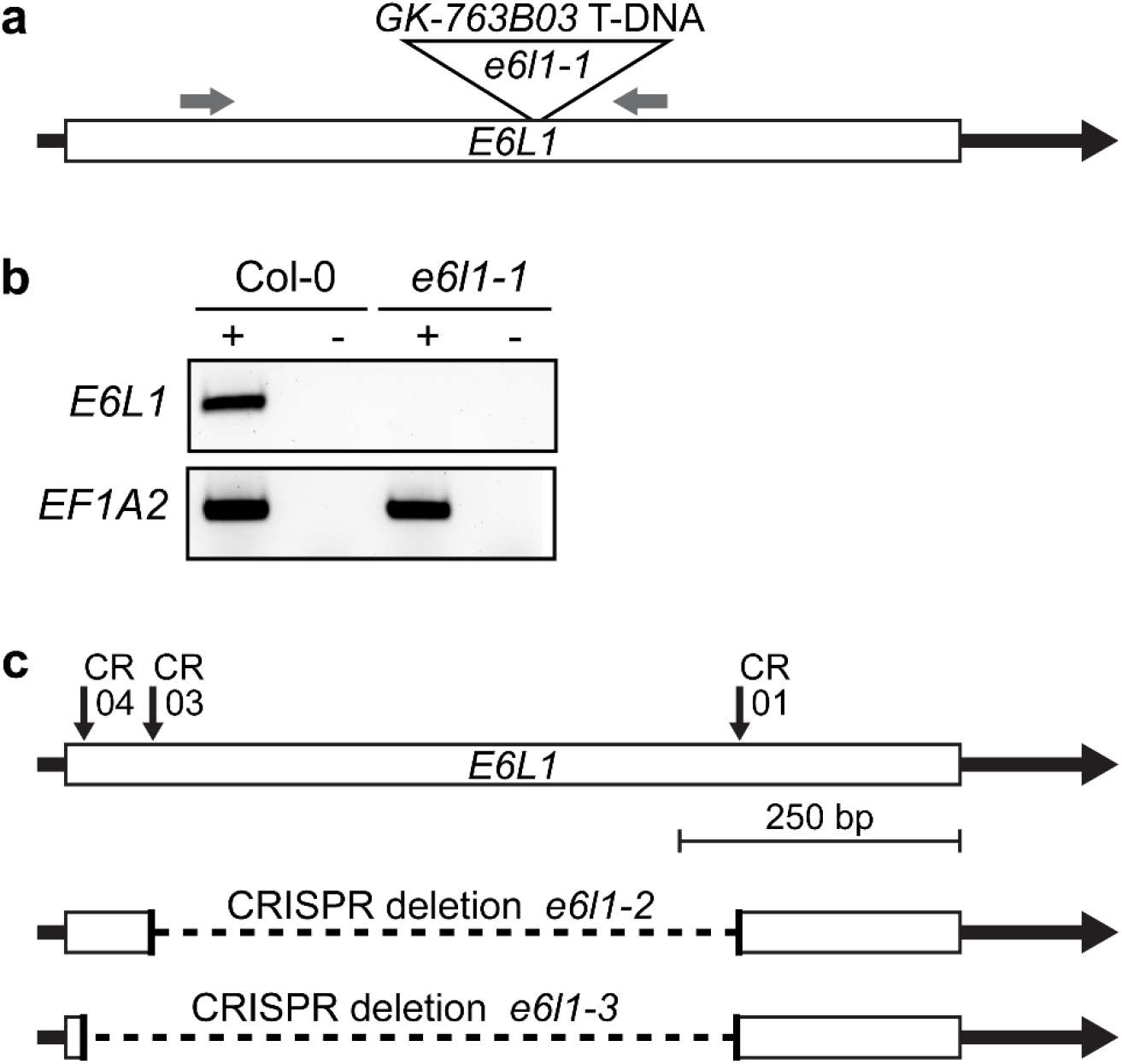
T-DNA and CRISPR sgRNA target locations for the *e6l1* mutants. **a.** *E6L1* gene structure (At2g33850) and T-DNA insertion site for the *e6l1-1* mutant (GK-763B03). Grey arrows on top are the primer locations for RT-PCR conducted in (**b**). **b.** RT-PCR analysis of *E6L1* expression in the *e6l1-1* mutant. The T-DNA successfully disrupted the *E6L1* gene as no expression was observed for the *e6l1-1* mutant when compared to wild-type Col-0. RT-PCR was performed using cDNA synthesized from stage 12 flower bud RNA. *EF1A2* (*ELF-1α*; At1g07930) was used as a positive control. *E6L1* PCR size = 430 bp and *EF1A2* PCR size = 350 bp. Reverse transcriptase was added (+) or not added (−) to each reaction, and the absence of any bands in the (−) lanes indicated that the absence of any genomic DNA contamination. **c.** CRISPR deletion mutations generated in the *E6L1* gene. Black arrows above the *E6L1* gene structure represent the sgRNAs used to generate the CRISPR deletions. The *e6l1-2* and *e6l1-3* mutants were generated using the CR01/CR03 and CR01/CR04 gRNA combinations, respectively.

Overall, the three homozygous *e6l1* mutants produced full-sized plants with flowers that were wild-type in appearance (Fig. 6A-D). However, upon closer inspection, two mild phenotypes related to the trichomes and stigmatic papillae were observed. For the trichomes, the *e6l1* mutant seedlings displayed wild-type branching with three-branched trichomes; however, the mutant trichome branches did not appear to be as straight as that observed for Col-0 (Fig. 7). Similarly, some of the stigmatic papillae displayed an increased curvature that was visible under both the light microscope (Fig. 6C-D) and SEM (Fig. 6G-J). The effect of this phenotype was that the *e6l1* stigmatic papillae looked a bit more disorganized (Fig. 6H-I) when compared to the wild-type Col-0 stigmas (Fig. 6G). To quantify the observed curvatures for the stigmatic papillae, measurements were taken from the SEM images of the angle of deviation of a papilla tip relative to its base. Stigmatic papillae from the three *e6l1* mutants displayed a significant increase in the deviation angle relative to Col-0 papillae (Fig 8). Given the mild stigmatic papilla phenotype, seed set rates in naturally pollinated siliques were also examined and a modest reduction in seed numbers was observed with an average of 46 seeds/silique for the three *e6l1* mutants in comparison to 52 seeds/silique for Col-0 (Fig. 9A). The source of this slight reduction appeared to due to some of the ovules not being fertilized as seen in dissected mature siliques (Fig. 10A-D). The position of the unfertilized ovules seemed to be random as seeds were also found at the base of the *e6l1* siliques (Fig. S4) and thus, not due to pollen tubes failing to reach the bottom of the transmitting tract.

**Fig. 6.**
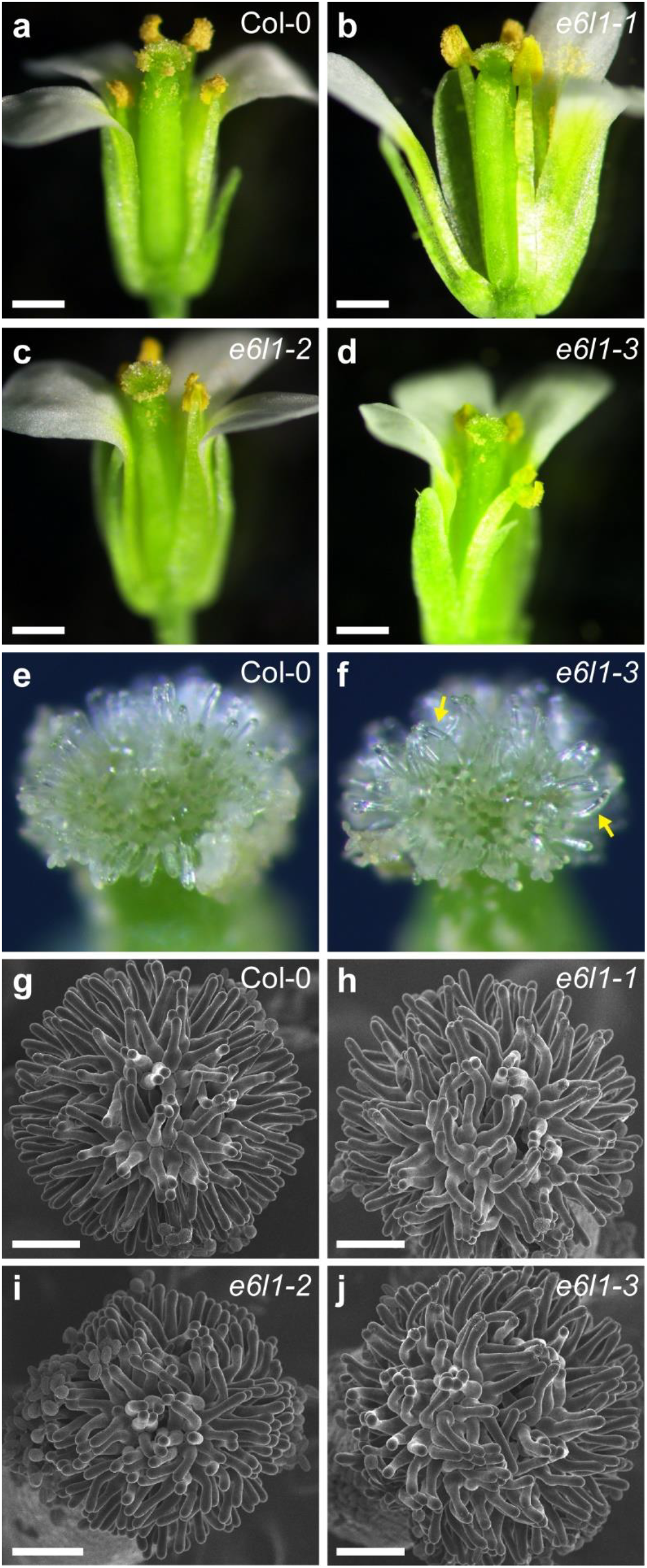
Representative images of stage 13 flowers and stigmas from the three *e6l1* mutants. **a-d.** Stage 13 flowers were dissected to show mature stigmas and anthers for Col-0 (**a**) and the three *e6l1* mutants (**b-d**). The *e6l1* mutant flowers were phenotypically normal in appearance. Scale bar = 500 μm. **e-f.** Light microscopy images of stigmas are shown for Col-0 (**e**) and the *e6l1-3* mutant (**f**). The *e6l1-3* mutant stigma displayed stigmatic papillae with a slight curvature (yellow arrow) when compared to Col-0. **g-j.** SEM images of stigmas are shown for Col-0 (**g**), e6l1-1 mutant (**h**), e6l1-2 mutant (**i**) and the e6l1-3 mutant (**j**). Many of the *e6l1* mutant stigmatic papillae are slightly curved in comparison to Col-0. Scale bar = 100μm.

**Fig. 7.**
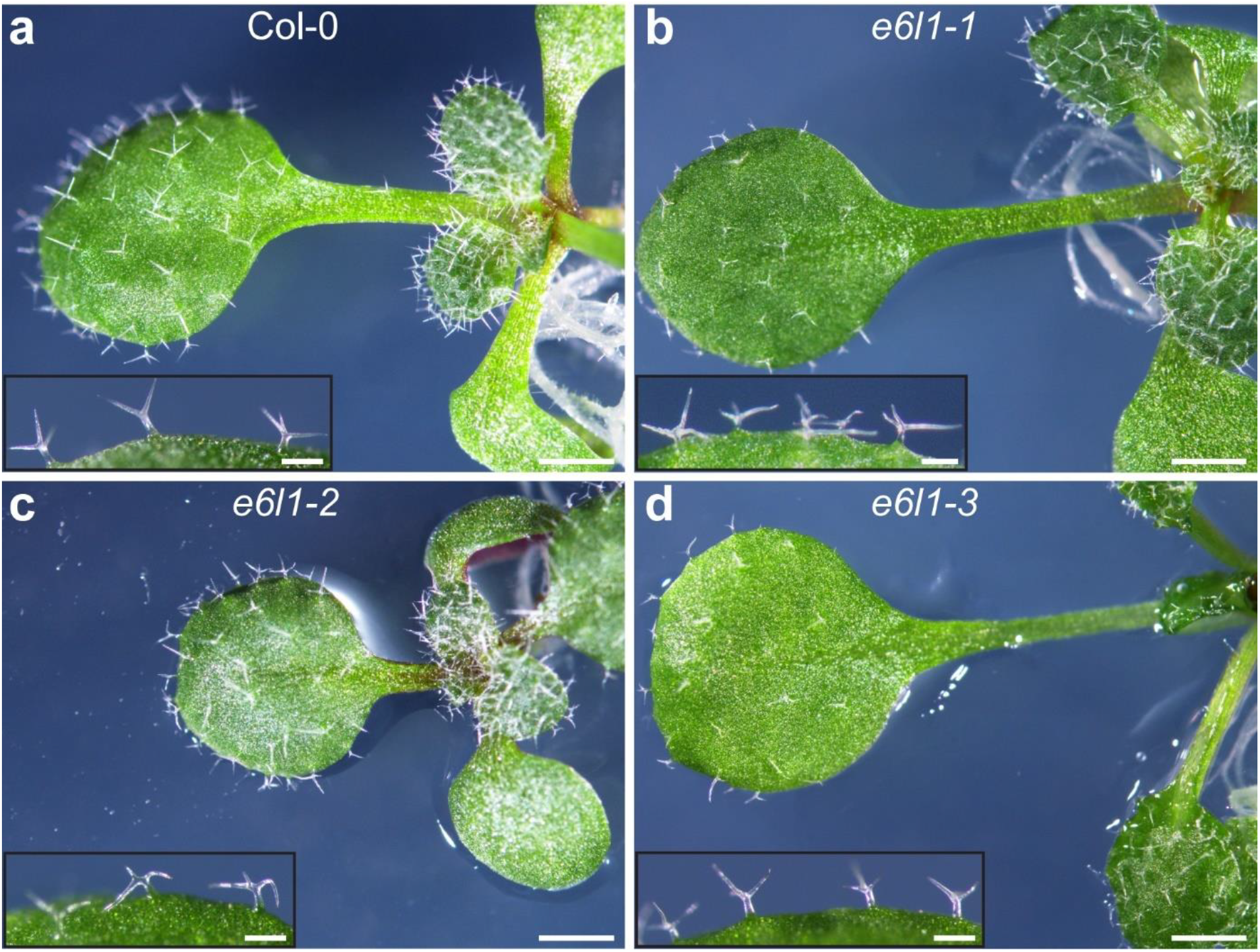
Representative images of trichomes on seedling leaves from the three e6l1 mutants. Trichome photos were taken on 2-week old seedlings grown on ½ MS plates with 1% sucrose. Scale bar = 1 mm for seedlings and 200 μm for the inset photos.

**Fig. 8.**
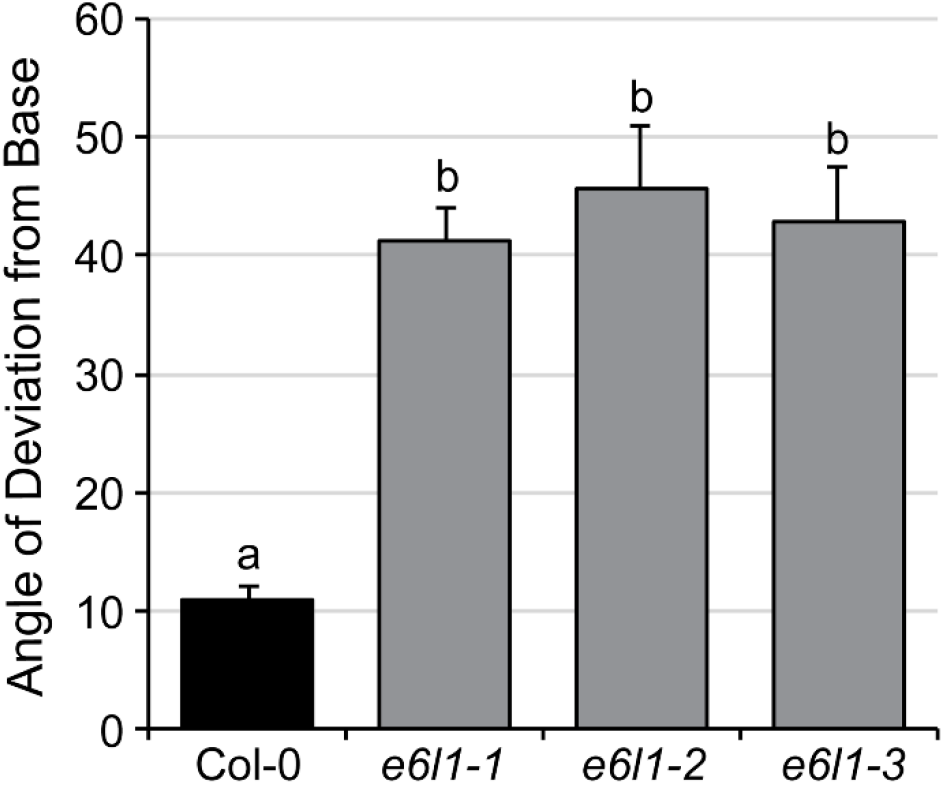
Angle of Deviation of the tip of the stigmatic papilla relative to the base. The observed curvatures of the stigmatic papillae for the *e6l1* mutant stigmatic papillae were quantified by measuring the angle of deviation of a papilla tip relative to its base in the SEM images. Measurements were taking using the Image J software (Schneider et al. 2012). n = 17 papillae for each sample. Letters represent statistically significant groupings of p<0.05 based on a one-way ANOVA with a Duncan post-hoc test.

**Fig. 9.**
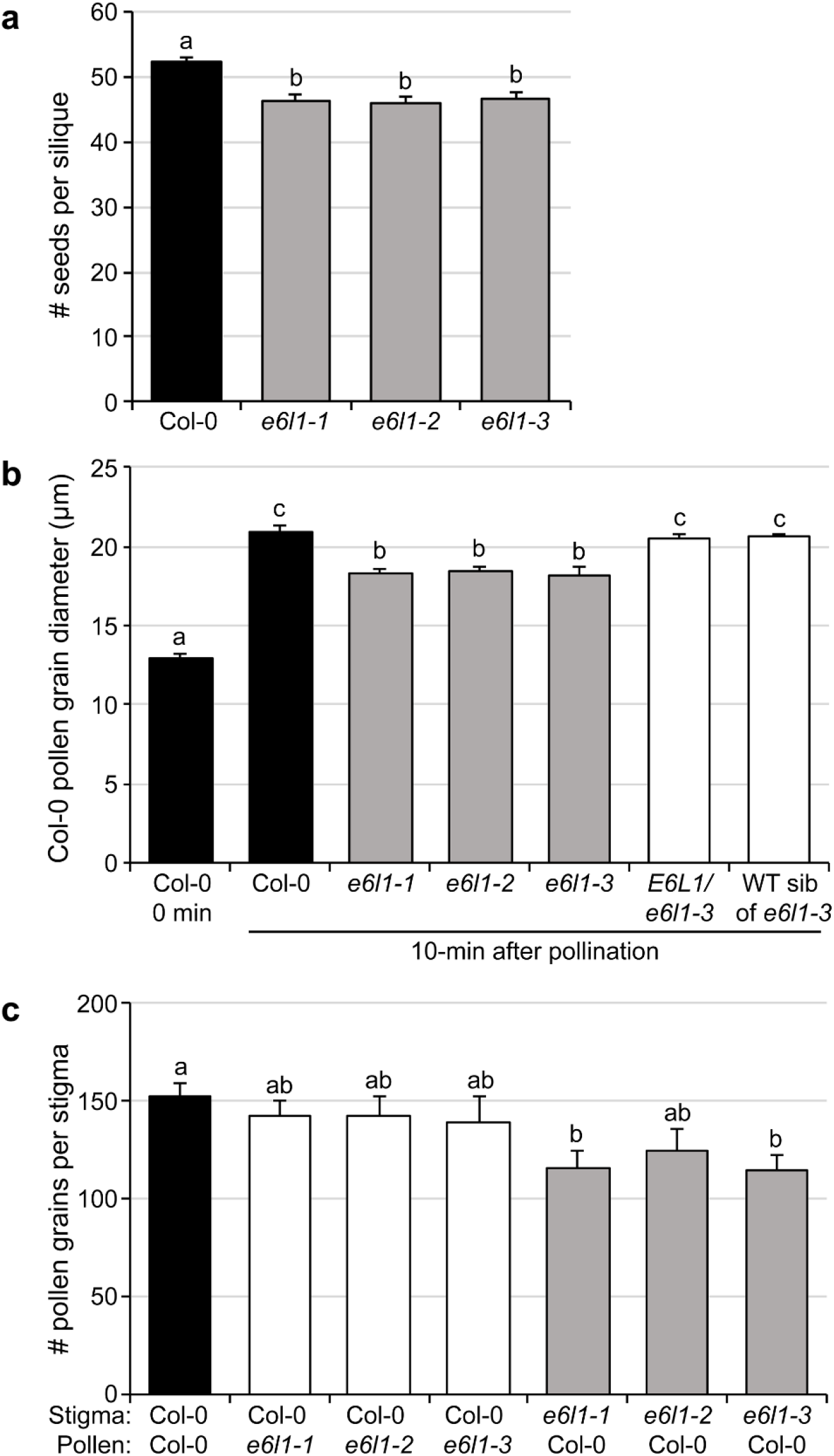
Graphs of post-pollination phenotypes for the three *e6l1* mutants. **a.** Seed set rates for naturally self-pollinated stigmas. Siliques from all three *e6l1* mutants showed a slight but significant reduction in seed set relative to Col-0; n = 10 siliques/line. **b.** Col-0 pollen hydration at 10 minutes post-pollination with Col-0 and *e6l1* mutant stigmas. In comparison to Col-0 stigmas, stigmas from the three *e6l1* mutants supported significantly reduced Col-0 pollen hydration. Results from a wild-type and *E6L1/e6l1-3* heterozygous sib of *e6l1-3* were wild-type as expected. Water uptake by the Col-0 pollen grains was measured using pollen grain diameter as a proxy; n = 30 pollen grains per line. **c.** Col-0 pollen grains adhered to stigmas following aniline blue staining at 2-hours post-pollination. All three *e6l1* mutants showed a small reduction in the number of Col-0 pollen grains remaining adhered to the aniline blue-stained samples when compared to Col-0 stigmas and the reciprocal crosses (Col-0 stigmas pollinated with *e6l1* mutant pollen); n = 10 stigmas for each cross. Letters represent statistically significant groupings of p<0.05 based on a one-way ANOVA with Duncan post-hoc tests for all samples.

**Fig. 10.**
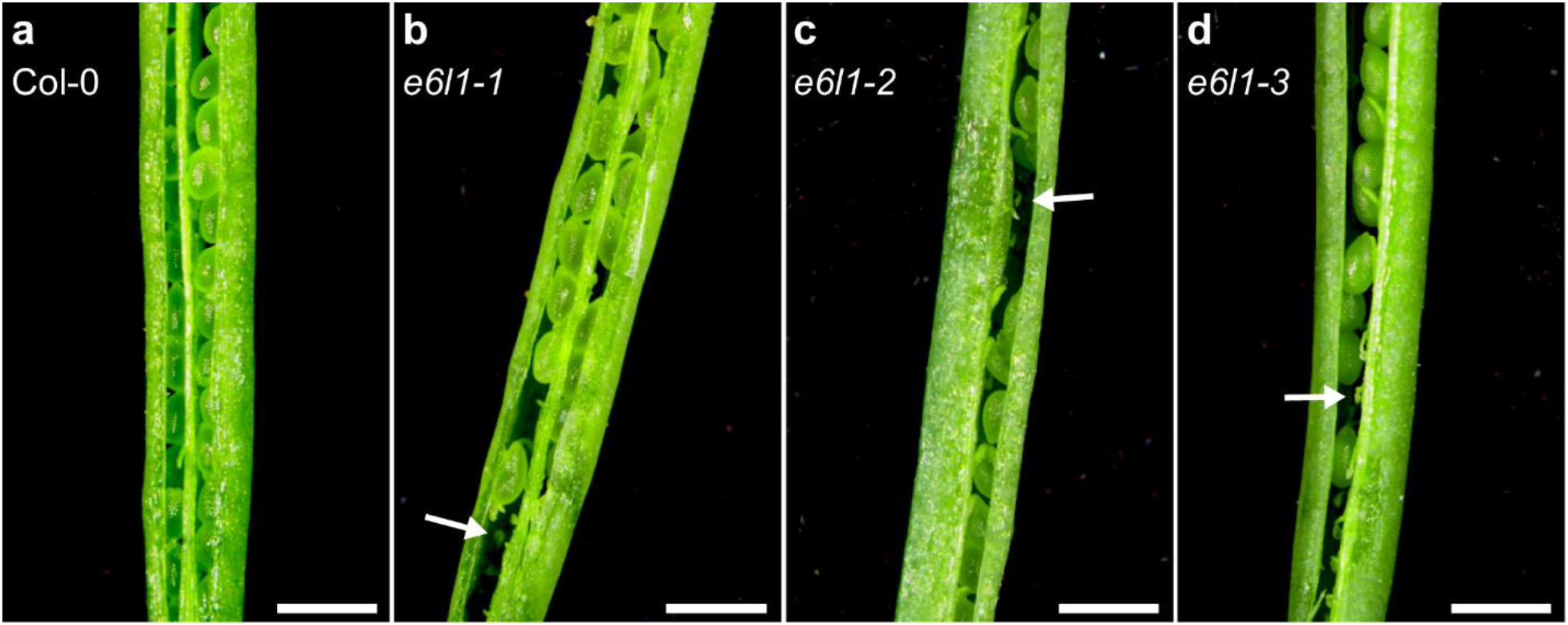
Representative images of fully developed opened siliques from the three *e6l1* mutants. **a-d.** Mature siliques were dissected longitudinally and opened. Full seed pods were observed for Col-0 (a), however some unfertilized ovules (white arrows) were observed for *e6l1* mutants (**b-d**). Scale bar = 1mm.

Given the high-level expression of E6L1 in the stigmatic papillae, the early stages of pollen-stigma interactions were scrutinized more closely in the three *e6l1* mutants. Following pollination, the *Arabidopsis* pollen grains require water from the stigma to rapidly hydrate and this step requires pollen recognition by the stigma (reviewed in (Doucet et al. 2016). Wild-type Col-0 pollen grains were placed on all stigmas, and their rate of hydration was examined at 10 min post-pollination by measuring Col-0 pollen grain diameters. The stigmas of the three *e6l1* mutants supported a significant reduction in Col-0 pollen hydration (~18μm) at 10 min post-pollination when compared to Col-0 stigmas (~21μm). For the *e6l1-3* mutant, a wild-type and *E6L1/e6l1-3* heterozygous sib was also tested, and Col-0 pollen displayed wild-type levels of hydration on these stigmas as expected (Fig. 9B).

Pollen tube growth through the stigma and the transmitting tract was examined by aniline blue staining of pollinated pistils. No visible differences were observed for pollen tube growth in naturally pollinated pistils (Fig. S5). We then examined this more closely using reciprocal pollinations between wild-type Col-0 and the three *e6l1* mutants. Again, there were no noticeable differences observed for the *e6l1* mutant pistils when the compared to Col-0 (Fig. 11A-H). The manually-pollinated aniline blue-stained pistils were also used to quantify the number of adhered pollen grains. These represent the well-adhered pollen grains as the remainder will have washed away during the staining process. There were no statistically significant differences between the Col-0 pistil samples pollinated with pollen from Col-0 and the three *e6l1* mutants as expected (Fig. 9C). However, when the *e6l1* mutant pistils were pollinated with Col-0 pollen, all three mutant lines displayed a reduction in the number of adhered pollen grains with averages ranging from 114 to 124 pollen grains/stigma compared to an average of 153 pollen grains/stigma for Col-0, and these reductions were statistically significant for the *e6l1-1* and *e6l1-3* pistils (Fig. 9C). In conclusion, three independent *e6l1* mutants showed reduced rates of wild-type pollen hydration and pollen adhesion, supporting our hypothesis that the stigma-expressed *E6L1* plays a role in stigma for pollen-stigma interactions.

**Fig. 11.**
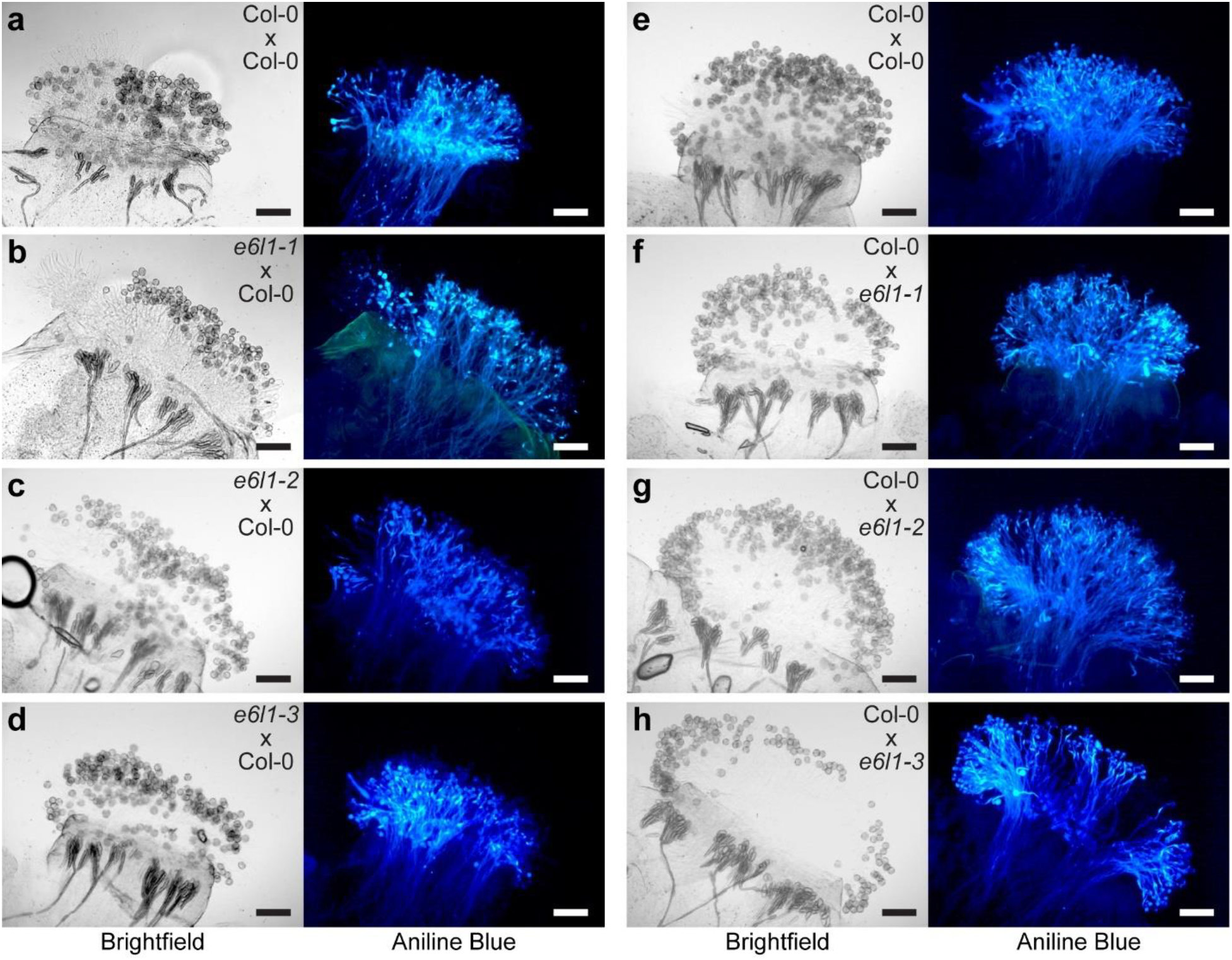
Representative images of aniline blue-stained stigmas from the three *e6l1* mutants. Reciprocal crosses were performed, and pollinated pistils were collected at 2 hours post-pollination for aniline blue-staining. Pollen grains adhered to the stigmatic papillae are shown in the brightfield images while aniline blue-stained pollen tubes are visible in the fluorescent images. Indicated crosses = female × male. **a-d.** Col-0 pollen grains were placed on Col-0, *e6l1-1*, *e6l1-2* and *e6l1-3* stigmas, respectively. **e-h.** Col-0, *e6l1-1*, *e6l1-2* and *e6l1-3* pollen grains, respectively, were placed on Col-0 stigmas. All crosses showed large numbers of adhered pollen grains and pollen tube growth. Scale bar = 100μm.

## Discussion

In this study, we have examined the function of the novel *Arabidopsis E6L1* gene in the stigma during early post-pollination processes. *E6L1* was of particular interest as it was one of the top-ranked genes based on expression levels in an *Arabidopsis* stigmatic papillae RNA-Seq dataset (Gao et al. 2018). The E6L1 protein was predicted to contain a signal peptide for secretion, and this was confirmed when an E6L1:RFP fusion was found to be localized to the apoplastic space in *N. benthamiana* epidermal cells. Since *E6L1* is highly expressed in the receptive stigmatic papillae, it is likely already secreted to the stigmatic papillar cell wall prior to pollination. Furthermore, the analysis of three independent *e6l1* mutants, including one T-DNA lines and two CRISPR deletion lines, uncovered mild fertility defects. Post-pollination defects included reduced wild-type pollen hydration and adhered pollen grains on the mutant *e6l1* stigmas, supporting a requirement for E6L1 in the early stages of this process.

In cotton, it was first hypothesized that the E6 protein may be a structural component of the cell wall in cotton fibers, but no conclusive evidence was presented to support this (John 1996; John and Crow 1992). It may be that the cotton E6 antisense suppression construct used by John (1996), did not sufficiently knockdown cotton E6 and closely-related *E6-like* members in this allotetraploid species to uncover a phenotype. Cotton fibers are single cell trichomes on the ovule surface, and interestingly, *Arabidopsis E6L1* was also found to be highly expressed in the leaf trichome microarray datasets. While no alterations to trichome branching were observed for the *e6l1* mutant leaf trichomes, we did notice that the branches were not always straight suggesting that E6L1 is structurally required to produce the three straight trichome branches seen in wild-type trichomes. The mild curvature observed for stigmatic papillae in the mutant *e6l1* stigmas would also be consistent with this proposed structural function.

A number of other genes have been found to display enriched expression in the elongation phase of cotton fiber development, including genes associated with cell elongation and cell wall loosening (Chen and Burke 2015; Ji et al. 2003; Lee et al. 2006; Lee et al. 2007; Li et al. 2002). One gene of particular note was the cotton orthologue of the *Arabidopsis FIDDLEHEAD (FDH)* gene which was found to be highly expressed in the developing cotton fibers (Lee et al. 2007; Li et al. 2002). As previously mentioned, the *Arabidopsis fdh* mutant has altered surface cuticles which then allowed for promiscuous germination of wild-type pollen on non-reproductive plant tissues (Lolle and Cheung 1993; Voisin et al. 2009). This observation ties back to the different categories of players implicated in early pollen-stigma interactions, and where E6L1 might fit in (‘structural’, ‘perception’ and ‘cellular responses’; as described in the introduction). Like FDH, E6L1 appears to play a more structural function in the *Arabidopsis* stigmatic papillae and trichomes, specifically in the cell wall where it would be localized. In addition, like other players implicated in defining structural features for the pollen or stigma (e.g. CER, FDH, OFT1), loss-of-function *e6l1* mutations also impact compatible pollen-stigma interactions.

Overall, E6-like proteins are broadly conserved, but restricted to flowering plant species. As part of our investigations into this protein family, we identified a 20 amino-acid region that is highly conserved in E6-like protein homologues. This domain is presumably related to their function in the cell wall as sequence searches using a consensus sequence for this region only identified E6-like homologues. Furthermore, this suggests that cotton E6 and E6-like proteins are neither uniquely involved in the process of cotton fiber development, nor Brassicaceae-specific pollen-stigma interactions. It is therefore possible that E6 proteins play a more generalized role in the cell wall, and are deposited during cell elongation processes, which would be consistent with both cotton fiber elongation and stigmatic papillar elongation. Our phylogenetic analysis of a subset of E6-like proteins did produce four supported family-specific clades and perhaps, these are connected to specific biological functions. Two of these clades include all the Brassicaceae E6-like protein homologues. It would be interesting, in particular, to investigate the expression patterns of the other Brassicaceae members in the *Arabidopsis* E6L1 clade to determine if they also display stigma-specific expression and a related function. Overall, the current work suggests that E6L1 is an important structural component in the stigmatic papillae and is required for the early stages of pollen-stigma interactions in *Arabidopsis*.

## Author contribution statement

JD and DRG designed the research. AD, ZG and MKN provided unpublished stigmatic papillae transcriptome datasets for further analysis. JD, CT, EFW and HKL performed the research. JD and DRG conducted the data analysis, prepared the figures and wrote the manuscript. JD, DRG and HKL edited the manuscript.

## Acknowledgements

We are very grateful to Alexander Leydon (Mark Johnson lab, Brown University) for providing valuable advice on using the CRISPR/Cas9 system, and Audrey Darabie for assistance with the SEM. We acknowledge the Salk Institute Genomic Analysis Laboratory (SIGnAL), GABI-Kat, and the Arabidopsis Biological Resource Center (ABRC) for providing the sequence-indexed Arabidopsis T-DNA insertion mutant, and Addgene and ABRC for providing the CRISPR/Cas9 vectors. We also thank Betty Geng for technical assistance with genotyping plants, and members of the Goring lab for critically reading this manuscript. JD and HKL were supported by Ontario Graduate Scholarships (OGS), and this research was supported by a grant from Natural Sciences and Engineering Research Council of Canada (NSERC) to DRG.

## Electronic Supplementary Material

**Fig. S1.**
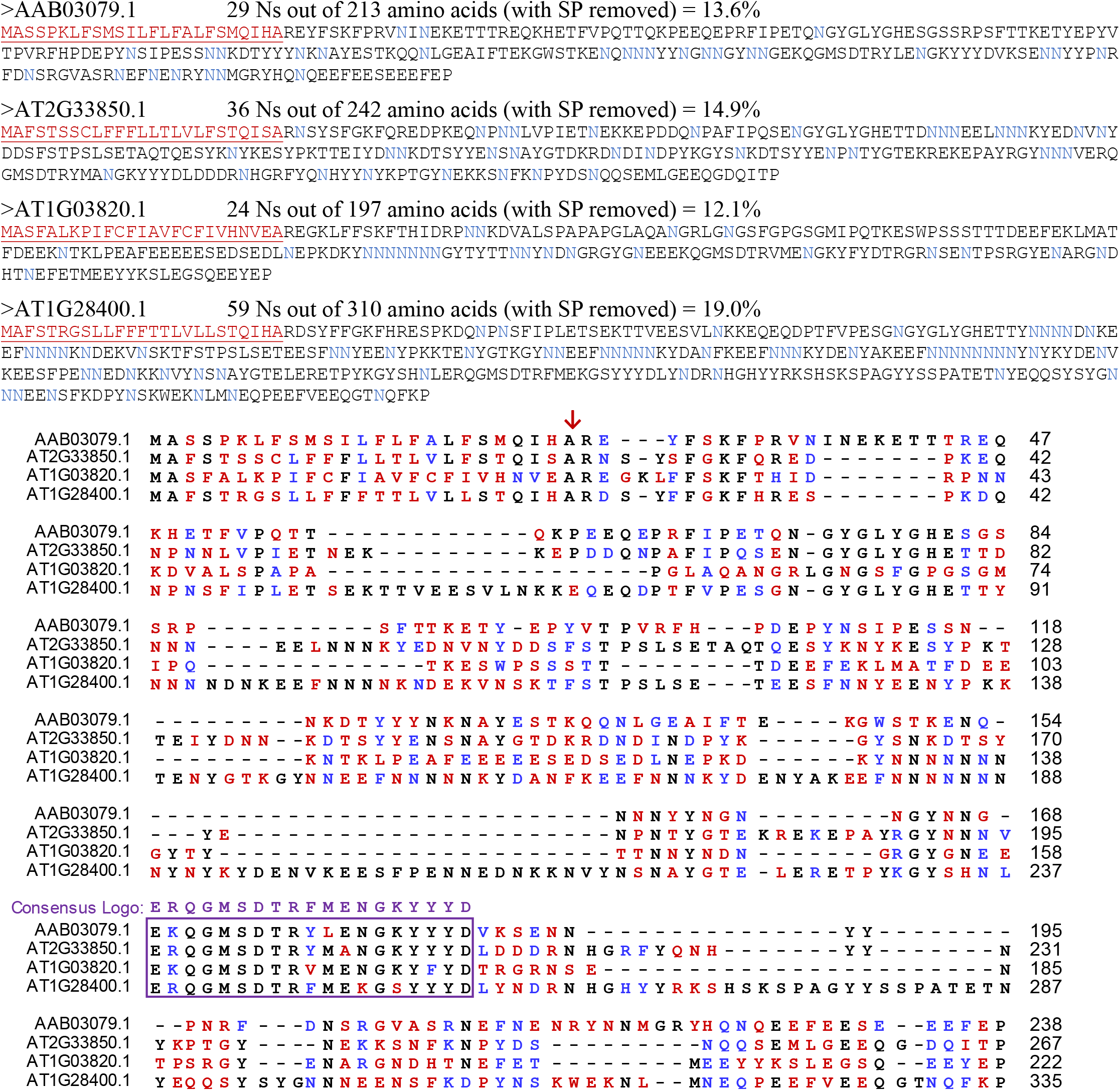
Amino acid sequence alignment of the predicted cotton E6 protein with three Arabidopsis homologues. The four sequences for the alignment are show at the top with the predicted signal peptides underlined in red (and cleavage site shown by red arrow in the alignment). The conserved E6 domain is boxed in the alignment with the Consensus Logo sequence shown above. Aligned sequences are: Cotton E6 protein (AAB03079.1) Arabidopsis E6L1 (AT2G33850.1), the second Arabidopsis ‘E6-like protein’(AT1G03820.1, Araport11), and the third Arabidopsis protein misannotated as a *GATA zinc finger protein* (AT1G28400.1, Araport11). The alignment of the four sequences was generated using Muscle in in MEGA7.0, and the alignment was formatted using Multiple Align Show (http://www.bioinformatics.org/sms/multi_align.html). Identical amino acids are shown in black text, similar amino acids are shown in blue text, and no identity are shown in red text. Percentage of sequences that must agree for identity or similarity coloring was set at 60%. Similar amino acids were grouped as: GAVLI (aliphatic), FYW (aromatic), CM (sulfur-containing), ST (hydroxyl), KRH (basic), DENQ (acidic & their amides), and P (cyclic) (http://www.biology.arizona.edu/biochemistry/problem_sets/aa/aa.html).

**Fig. S2.**
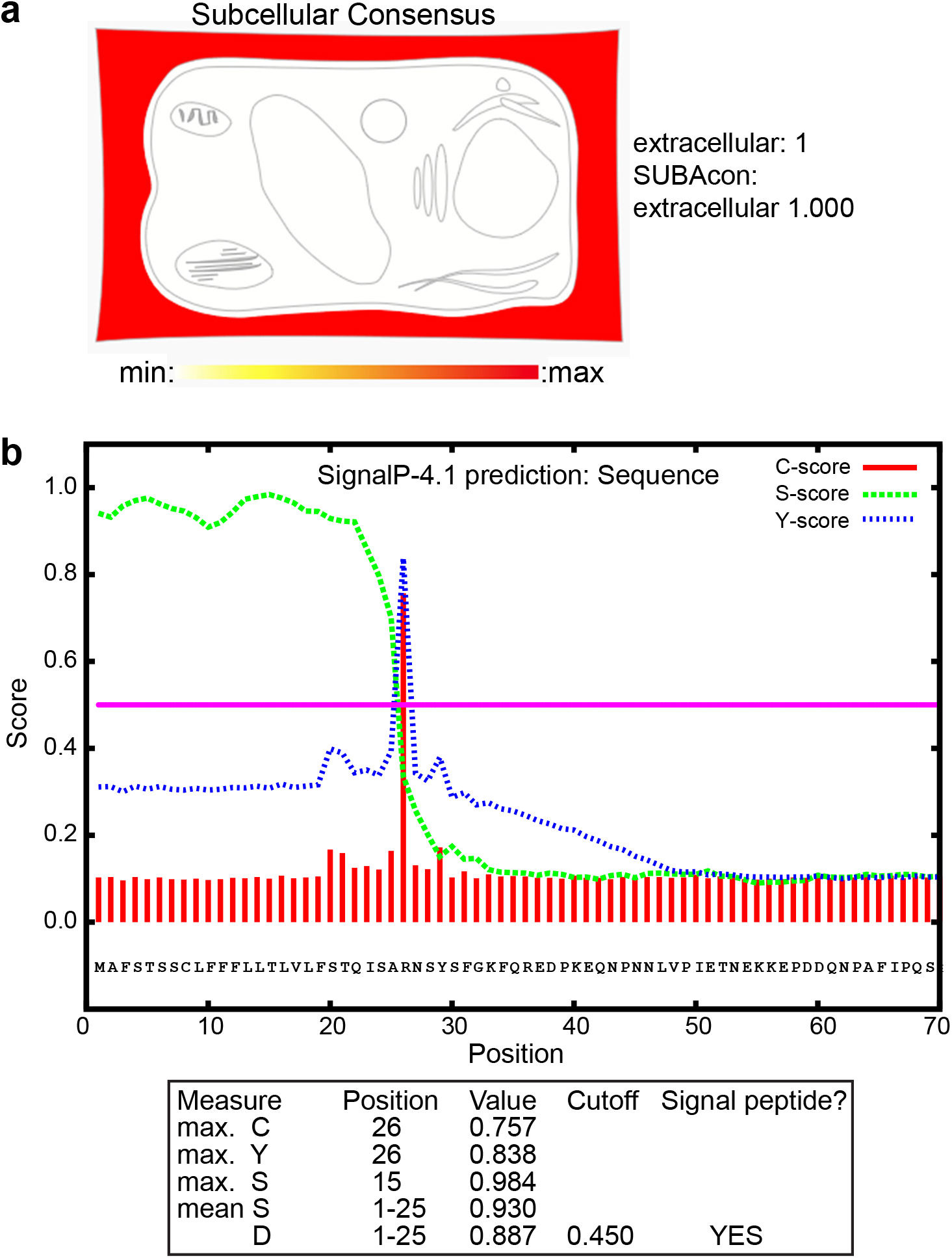
Predicted localization for E6L1 protein. a Subcellular localization prediction for E6L1 from SUBA4 SUBAcon shows predominantly extracellular localization based on prediction (Hooper et al., 2014; Hooper et al., 2017). b SignalP prediction for signal transit peptide. E6L1 is predicted to have a signal peptide prediction whereby the C-score represents the predicted cleavage position (red), the S-score represents the distinction between signal peptides from positions of mature proteins from proteins without signal peptides (green), and the Y-score represents the combined average of the two scores (blue). The cleavage site of the signal peptide is predicted to be at position 25 in the amino acid sequence (Petersen et al., 2011).

**Fig. S3.**
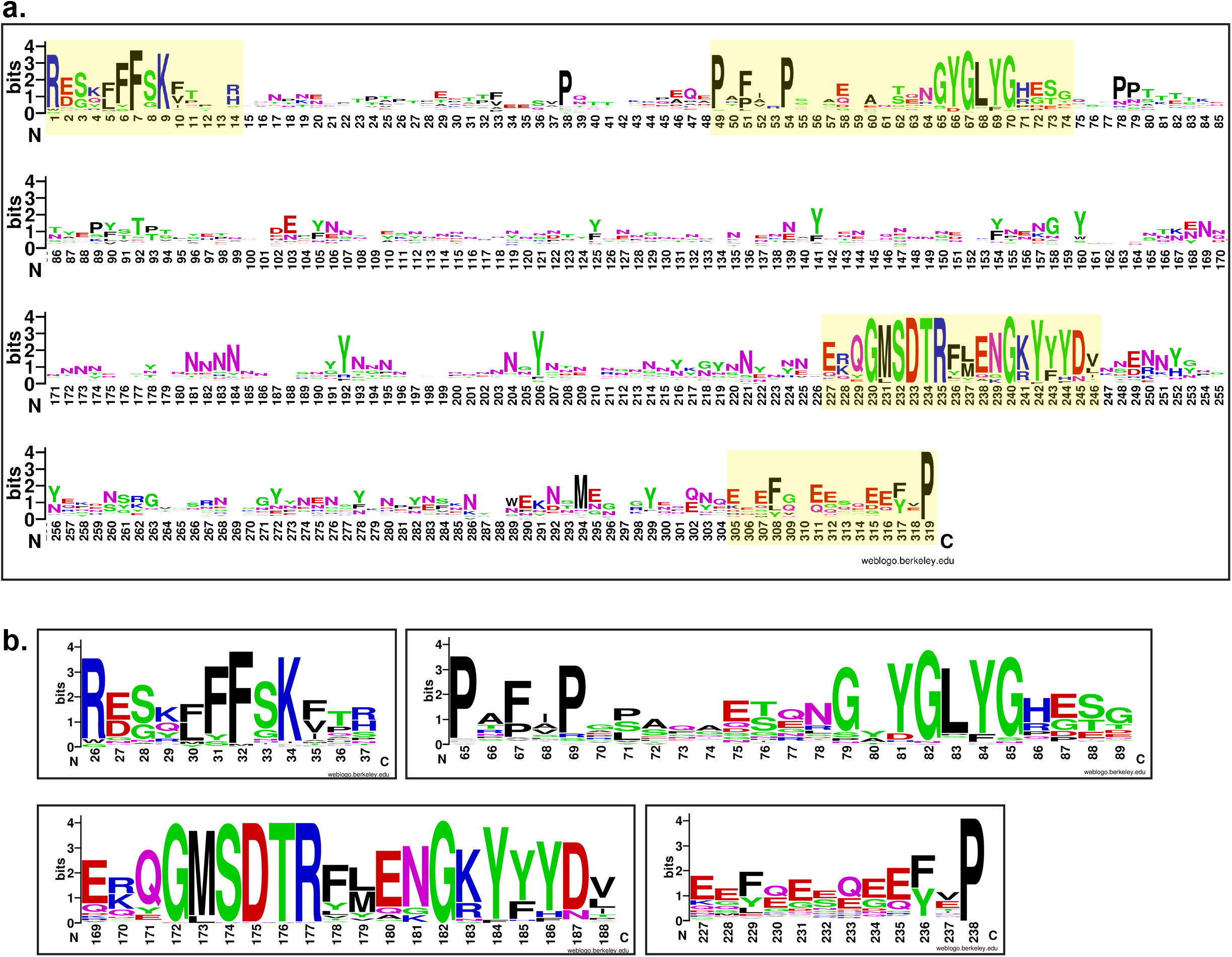
Consensus Sequence Logos for the predicted Angiosperm E6-like proteins. **a.** A Consensus Sequence Logo for the entire E6-like protein alignment. An alignment of 73 E6-like protein sequences was generated using Muscle (Edgar 2004) in the MEGA 7 software (Kumar et al. 2016). Signal peptides and large gaps in the alignment were removed prior to generating the full-length consensus sequence logo using the WEBLOGO tool (Crooks et al. 2004). The numbering on the X-axis starts at the first amino acid after the signal peptide. See File S3 for the alignment. **b.** Consensus Sequence Logos for four conserved regions. The four regions showing conservation are highlighted in **a**, and each region was specifically re-aligned to generate new Consensus Sequence Logos. Numbers for these corresponds to positions in the cotton E6 amino acid sequence. See Files S4-S7 for alignments.

**Fig. S4.**
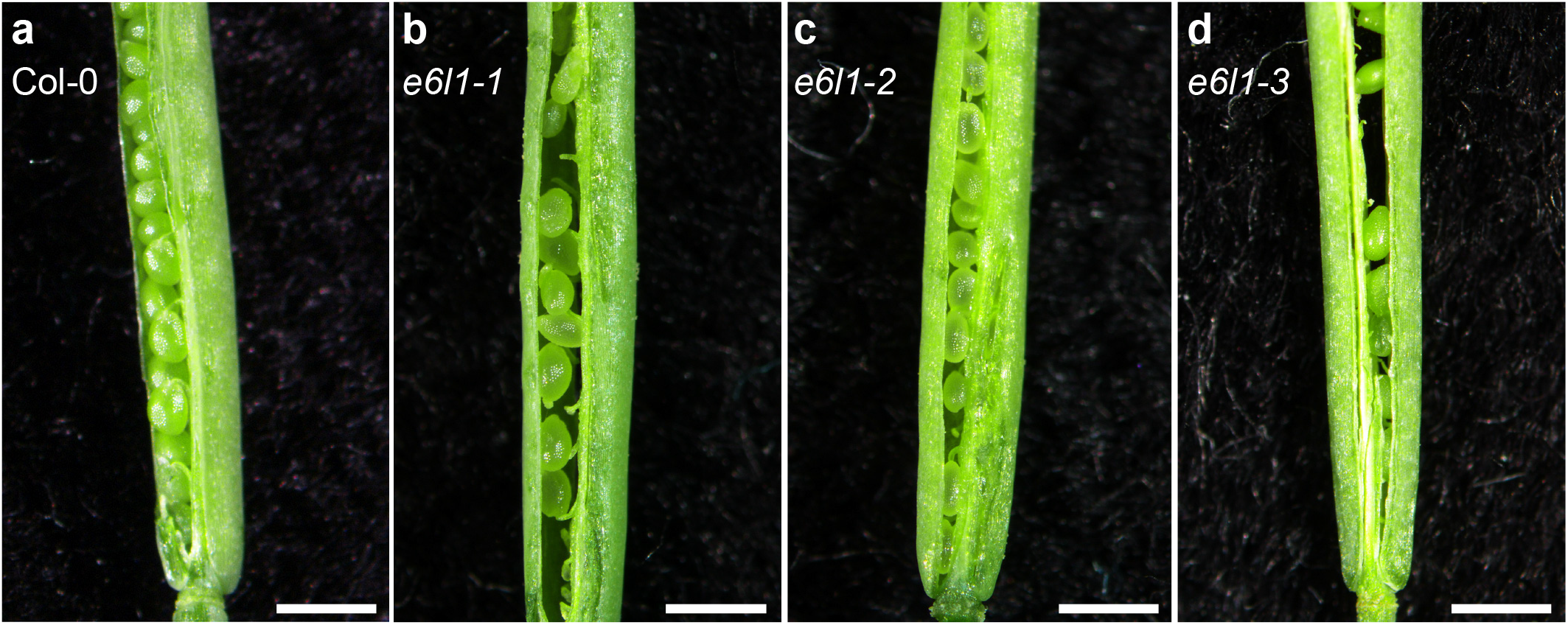
Representative images at the base of fully developed opened siliques from the three *e6l1* mutants. These images cover the base of the silique showing that there are seeds present in some instances. There was no consistent pattern indicating that the position of the seeds was random and that pollen tubes were able to grow to the base. Scale bar = 1mm

**Fig. S5.**
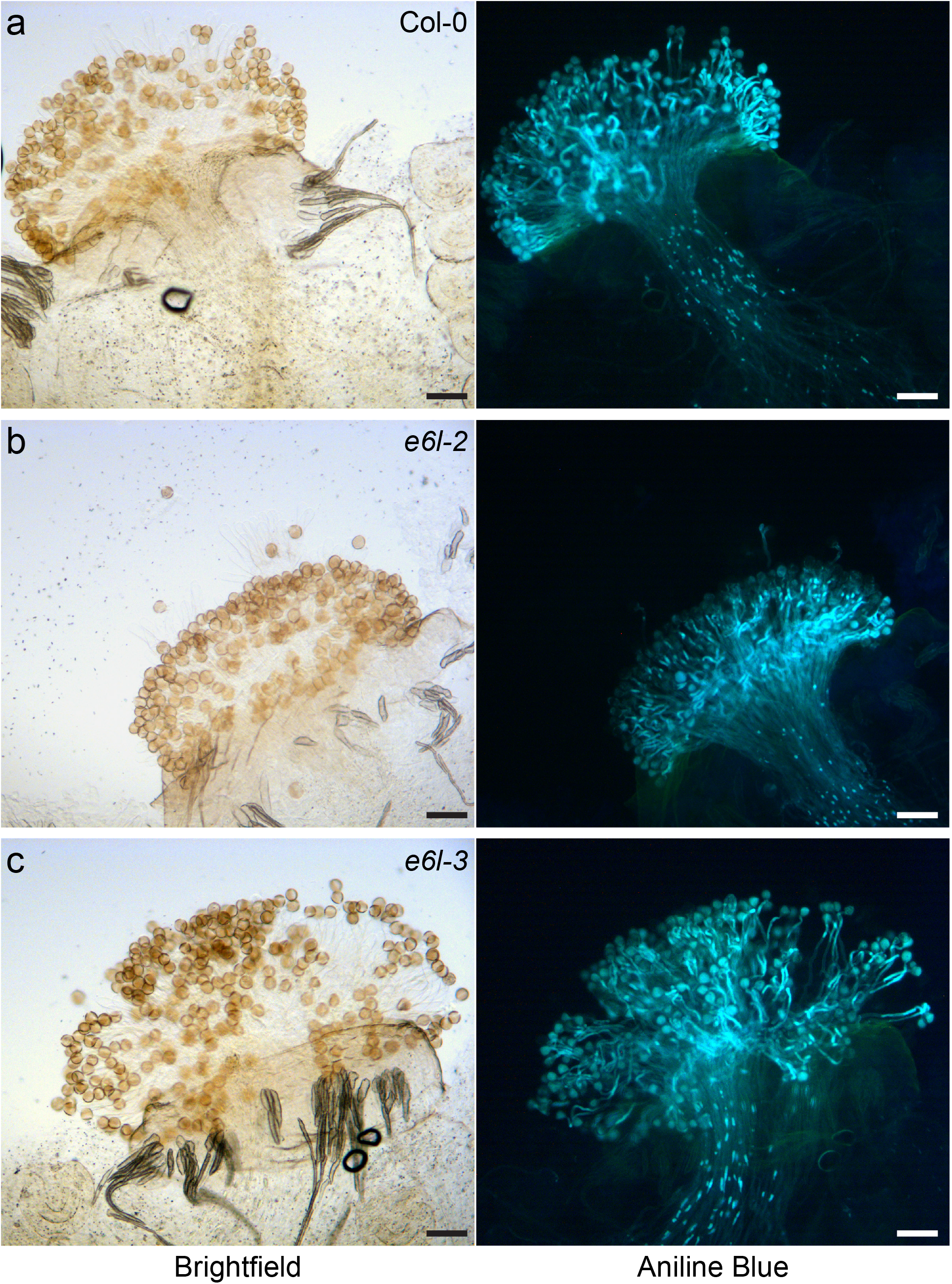
Representative images of aniline blue-stained stigmas from naturally pollinated pistils. Open flowers were inspected for pistils that had been already pollinated, and these pistils were collected for aniline blue staining. Pollen grains adhered to the stigmatic papillae are shown in the brightfield images while aniline blue-stained pollen tubes are visible in the fluorescent images. Scale bar = 100μm

**Table S1** BAR Expression Angler with SLR1 (At3g12000) as Bait (Top 25 hits).

**Table S2** BAR Expression Angler with SLR1 (At3g12000) as Bait (Top 25 hits): Subset of Tissues queried in Expression angler.

**Table S3** Top 25 Expressed Genes in the Stigmatic papillae RNA-Seq datasets.

**Table S4** Top 25 Expressed Genes in the Stigma Microarray datasets.

**Table S5** TRAVA RNA-Seq Datasets.

**Table S6** Primers used in this study.

**File S1** 73 Angiosperm E6-like amino acid sequences

**File S2** Alignment of 73 Angiosperm E6-like sequences for Fig. 4

**File S3** Alignment of Angiosperm E6-like sequences for Fig. 3 and Fig. S3.

**File S4** Alignment of E6-like conserved domain-26 for Fig. S3.

**File S5** Alignment of E6-like conserved domain-65 for Fig. S3.

**File S6** Alignment of E6-like conserved domain-169 for Fig. 3 and Fig. S3.

**File S7** Alignment of E6-like conserved domain-227 for Fig. S3.

